# Developmental Dynamics of Green Fluorescent Chromatophores in the Daggerblade Grass Shrimp, Palaemonetes Pugio Holthuis, 1949 (Decapoda, Caridea, Palaemonidae)

**DOI:** 10.1101/396648

**Authors:** M. P. Phelps

## Abstract

The daggerblade grass shrimp, *Palaemonetes pugio* Holthuis 1949 relies heavily on transparency as the primary form of camouflage yet possess several types of pigmented chromatophores located throughout the body. A distinct sub-population of yellow/white chromatophores have been discovered to exhibit brilliant green fluorescence. These cells develop in the embryo and are the primary chromatophore present in larval organisms. Post-larval grass shrimp undergo a major restructuring of the pattern and morphology of fluorescent chromatophores after metamorphosis with chromatophores found uniformly distributed throughout the body and at high concentration on the hepatopancreas and the eye stalks. In adult *P. pugio* the number of fluorescent chromatophores is significantly reduced and fluorescence is limited to only a subset of these chromatophores. The novel fluorescent properties of these cells, there relatively high abundance during early life stages, and pattern of development, suggest important cellular functions for these fluorescent chromatophores in grass shrimp.

## INTRODUCTION

Daggerblade grass shrimp, *Palaemonetes pugio* Holthuis, 1949, are widespread across the North American Atlantic coast ranging from the Gulf of Mexico to Nova Scotia (Berg and Sandifer, 1984). In this region *P. pugio* is one of the most abundant macro invertebrates inhabiting shallow water salt marshes and estuarine environments (Anderson, 1985). A key evolutionary adaptation for *P. pugio* is its whole-body transparency that is maintained throughout all larval to adult life stages. While transparency is a highly effective form of camouflage, development of transparent tissue often requires physiological tradeoffs (Johnsen, 2001; Bhandiwad and Johnsen, 2011; Bagge et al., 2017). Transparent organisms can also be susceptible to the harmful effects of ultraviolet (UV) radiation since pigmentation plays an important role as a UV protectant (Johnsen and Widder, 2001; Hansson, 2004; Hansson, Hylander, and Sommaruga, 2007).

Despite being highly transparent *P. pugio* possess several types of pigmented chromatophores as both larvae or adults (Brown, 1933). The chromatophores of *Palaemonetes spp.* have been studied extensively in the past with regard to the factors involved in pigment migration in crustaceans (Perkins and Snook, 1932; Brown, 1935; Brown, Webb, and Sandeen, 1952; Fingerman and Tinkle, 1956; Lambert and Fingerman, 1978, 1979). The first of these chromatophores develop in the embryo and are easily observed through the transparent chorion (Glas et al., 1997, Romney and Reiber, 2013). It has been noted that these embryonic chromatophores exhibit green fluorescence (Glas et al., 1997), which has been attributed to autofluorescence, however the development and prevalence of these fluorescent chromatophores in larval, juvenile and adult grass shrimp is unknown.

Cellular fluorescence can be derived from the accumulation of fluorescent pigment molecules or proteins. One of the most common fluorescent pigments are pteridine pigments, which are present in the chromatophores of a number of diverse organisms (Ortiz and Williams-Ashman, 1963; Stackhouse, 1966; Epperlein and Löfberg, 1984; Guyader and Jesuthasan, 2002). Fluorescent proteins are a novel adaptation most commonly found in Cnidarians (Chudakov et al., 2010). A growing number of marine organisms have been found to utilize fluorescent proteins including several species of copepod crustaceans and the lancelet, *Branchiostoma floridae* Hubbs, 1922 (Baumann et al., 2008; Bomati, Manning, and Deheyn, 2009; Hunt et al., 2010). The evolutionary benefits of many fluorescent adaptations however, are still unclear (Shagin et al., 2004; Chudakov et al., 2010).

This study details the developmental changes of green fluorescent chromatophores in *P. pugio* through embryonic, larval and adult live stages. Changes in cellular morphology and interaction with secondary chromatophores are analyzed to develop a map of chromatophore development in grass shrimp. This research also profiles the reduction of fluorescent chromatophores in adult shrimp and discusses potential functional implications of such pigmentation in the life history of marine grass shrimp.

## MATERIAL AND METHODS

### Animals Husbandry

Adult *P. pugio* were originally obtained from Midway Inlet, South Carolina, USA. Shrimp were communally housed in 38L aquariums with independent heating, aeration and filtration. All shrimp and developing embryos were cultured in 22-26 ppt artificial sea water (Instant Ocean, Blacksburg, VA) maintained at 25°C with a 15:9, light:dark photoperiod. Eggs were collected from anesthetized (0.3 ppt clove oil), gravid females using forceps and incubated on a rocking shaker until hatched. Newly hatched larvae were reared in 0.5 L plastic tubs in a temperature-controlled incubator with regulated photoperiod and daily water changes. Newly hatched Artemia nauplii were fed to all larvae once a day with post larvae and adults fed commercially formulated feed (Otohime) twice a day.

For time series analysis, larval shrimp were maintained individually in 12 well plates. Daily water changes were performed along with the daily feeding of Artemia nauplii. A total of 7 larvae were imaged at hatch and every other day, for a total of 8 days.

### Microscopy

Digital color images of live shrimp were taken throughout development with a high-resolution color camera (Digital Sight DS Ri1, Nikon, Japan), mounted to a Nikon SMZ1500 stereomicroscope. The microscope was equipped with blue (P-Blue), green (ET-GFP) and red (ET-DSRED) fluorescent filters (Nikon). Fluorescent excitation was provided by a 120W mercury vapor short arc light source. Images were acquired using NIS Elements software with exposure times ranging from 0.5 to 2 sec (Nikon).

All larvae were imaged without anesthesia by placing the animal in a drop of water on petri dishes coated with 1% agarose gel. While chromatophore development was followed throughout the larval cycle, detailed characterization of the ninth stage zoea (approximately 2 weeks post hatch) was used to determine the overall developmental changes in larval green fluorescent chromatophores prior to post larval metamorphosis. Anesthesia (0.3 ppt clove oil) was necessary for post-larval and adult shrimp, but all shrimp were imaged alive.

## RESULTS

### Green fluorescent chromatophores in larval *P. pugio*

A previous description of the embryonic development of *P. pugio*, noted fluorescent pigment cells in the late stage grass shrimp embryo (Glas et al., 1997). To validate the existence of these embryonic chromatophores and examine their role in larval pigmentation, late stage embryos and newly hatched larvae were examined with bright field and fluorescent microscopy (fig. 1). Green fluorescence was identified in the eyes and chromatophores of embryonic *P. pugio* (fig. 1C-C’), however the fluorescence in these cells was not due to tissue autofluorescence since it was specific to green emission wavelengths. These fluorescent chromatophores exhibited striking fluorescent intensity that was readily detectable with as little as 500 ms of exposure (fig. 1C-C’). The green chromatophore fluorescence was retained in all yellow/white chromatophores present in first stage zoea (fig. 1A-B). These chromatophores are the only distinct pigment cells observed in first zoea (Broad, 1957) and are dispersed into discrete clusters throughout the body (fig. 1A-B). Several small fluorescent chromatophores are located in the head region, at the base of the antennules and distributed on the anterior end of the carapace between the eyes (fig. 1D-D’). Dorsal green fluorescent chromatophores consist of a single large chromatophore located on the posterior margin of the second tergite (D3; fig. 1e-e’). Another cluster of paired fluorescent chromatophores reside on the posterior side of each eye (D1) and on either side of the base of the rostrum (D2; fig. 1 F-F’). Ventral fluorescent chromatophores consist of a pair of cells just posterior to the larval pereiopods and on abdominal sternites 2 and 3 (V1 and 2; fig. 1 G-G’). There is also a prominent fluorescent chromatophore located on the telson.

**Fig. 1.**
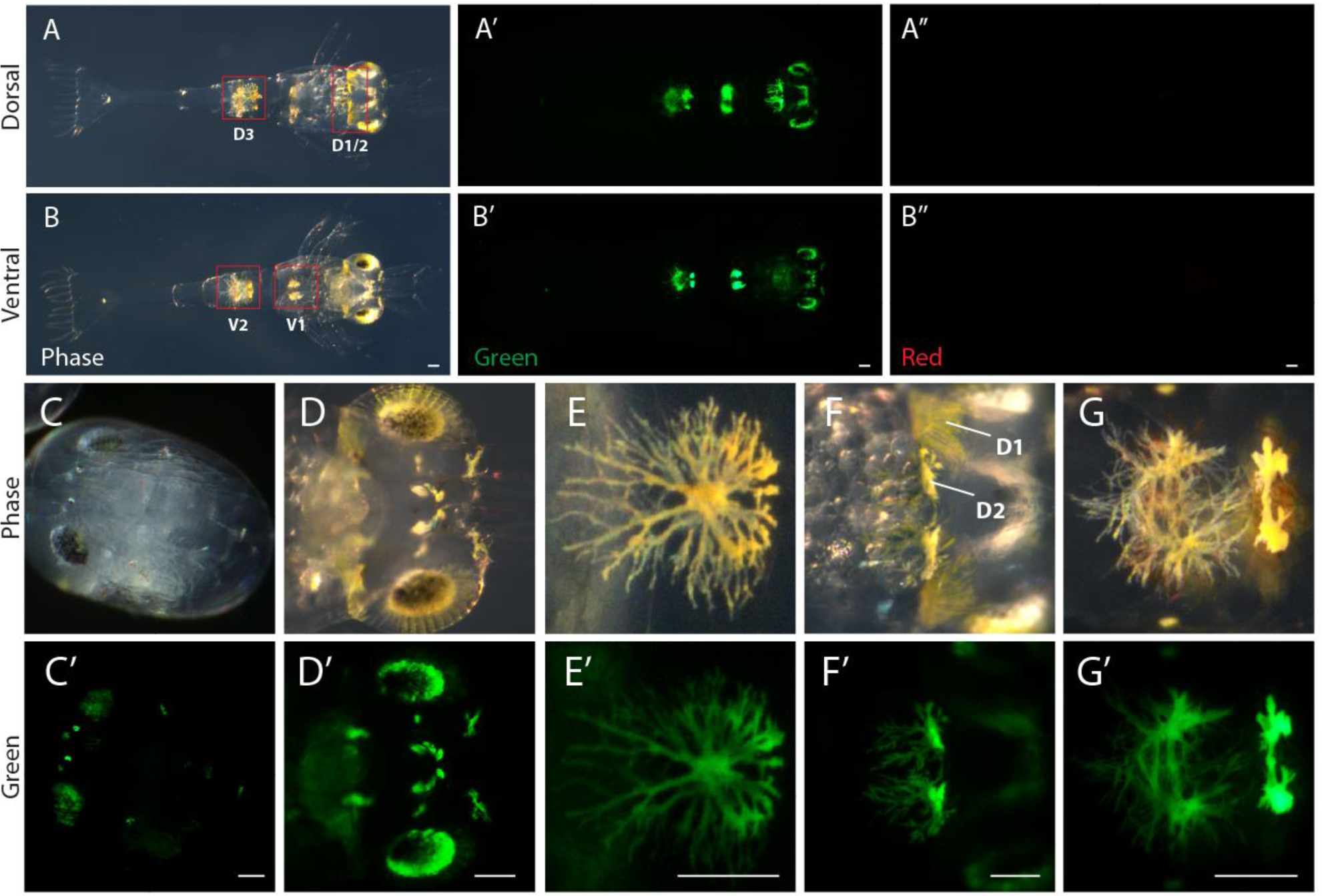
Green fluorescent chromatophores in the late stage embryo and first zoea of *P. pugio.* Whole body images of the dorsal (A-A”) and ventral surfaces (B-B”) of newly hatched first zoea exhibit green fluorescent chromatophores in the head region, including the eyes (D-D’), and throughout the body. These chromatophores are readily detectable in the late stage developing embryo (C-C’). Dorsal fluorescent chromatophores reside on the posterior end of the eyes (D1, F-F’), on either side of the rostrum (D2, F-F”), and on the second tergite (D3, E-E’). Pairs of ventral fluorescent chromatophores are located on the posterior end of the pereiopods (V1) and on abdominal sternites 2 and 3 (V2, G-G’). Background green fluorescence is caused by chromatophores out of the field of focus, which are visualized through the transparent body. Chromatophore clusters are identified by their anterior to posterior (1,2,3) location on either the dorsal (D) or ventral (V) surface. Images taken under phase contrast (Phase), and fluorescent green (’) and red (“) filters are identified. All scale bars represent 100μm

### Larval development of fluorescent chromatophores in *P. pugio*

Green fluorescent chromatophores are maintained throughout larval development. While there are no significant changes in fluorescent chromatophore number, there are small localized increases in the number of cells at several of the cluster locations. Cellular proliferation on the ventral abdominal chromatophore cluster (V2) and surrounding the telson give rise to a pair of additional green fluorescent chromatophores at these locations (fig. 2A-B). The chromatophores surrounding the telson appear to arise with the development of uropods since newly developed chromatophores are present at the base of each exopod (fig. 2a-a”). Proliferation of the solo large dorsal chromatophore (D3) also produces 2-3 additional fluorescent cells (fig. 2C-C’). The other chromatophore clusters did not appear to exhibit changes in cell numbers but morphological changes were evident (fig. 2D-F).

**Fig. 2.**
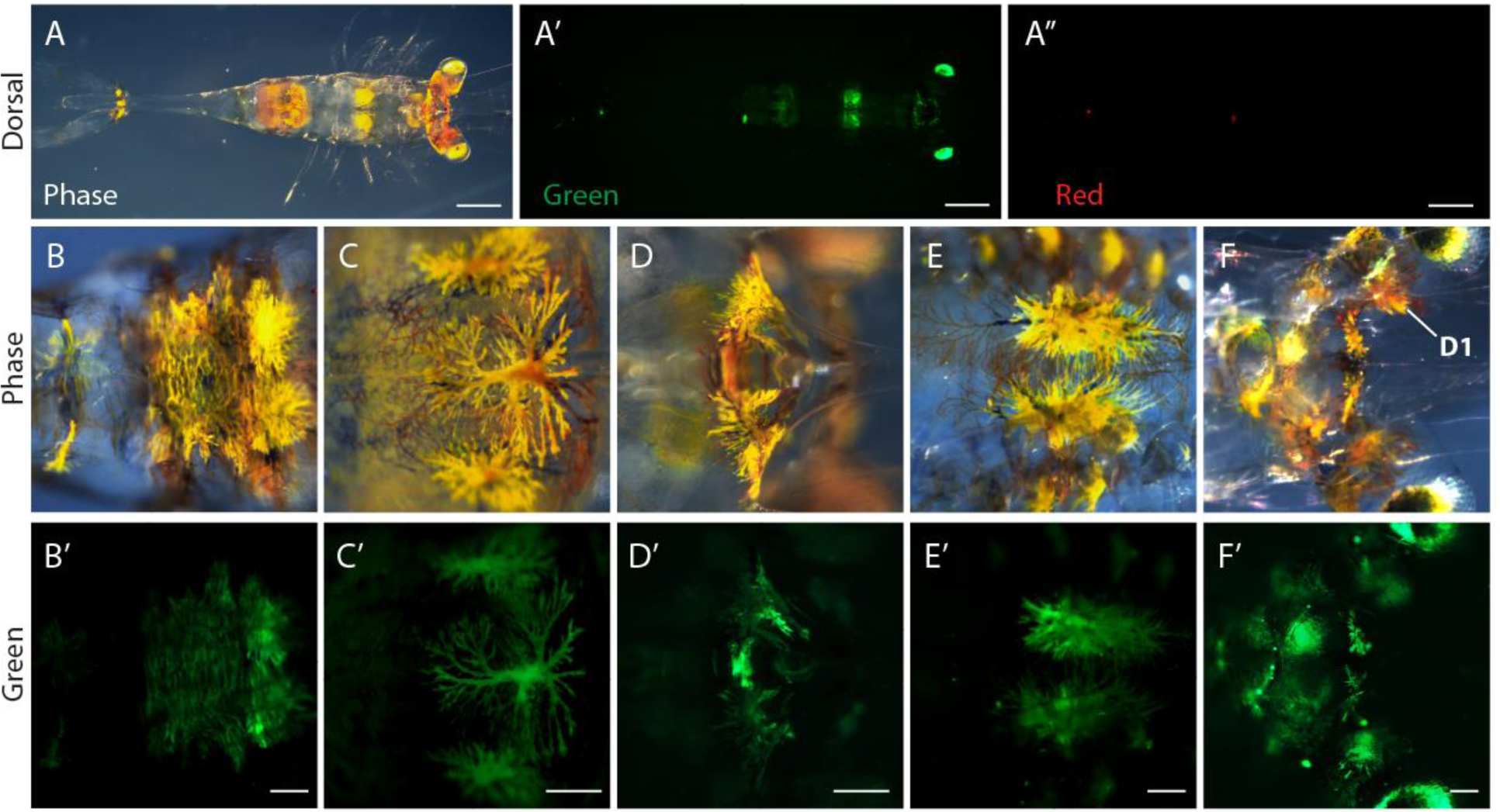
Pigmentation pattern in the ninth stage zoea, prior to metamorphosis. Dorsal whole-body images (A-A”) show little change in chromatophore locations compared to first zoea. The ventral, V2 cluster (B-B’) and the dorsal D3 cluster (C-C’) both develop additional fluorescent chromatophores. No change in chromatophore numbers were observed at cluster locations D2 (D-D’), V1 (E-E’), and on the head region, which includes cluster D1 located on the eye stalks (F-F’). Images taken under phase contrast (Phase), and fluorescent green (’) and red (“) filters are identified. Scale bars for whole body images (A-A”) and individual chromatophore clusters are 500 and 100μm, respectively. The observed background green fluorescence is indicative of chromatophores out of the field of focus, which are visualized through the transparent body.

One distinct change between first stage and ninth stage zoea is the co-localization of red and blue pigments on, or in close proximity to the fluorescent chromatophores. While interesting, the polychromatic nature of these pigment cells was not examined in this research since it has been studied previously in adult *P. pugio* (Brown, 1933, 1934). All larval green fluorescent chromatophore clusters exhibited these additional pigment cells which were non-fluorescent (fig. 2).

### Time series analysis of fluorescent chromatophore development

To gain insight into the recruitment of non-fluorescent chromatophores and their potential impact on the development of fluorescent chromatophores, time series analysis was performed by analyzing a single large dorsal chromatophore (D3) and surrounding tergites every other day for the first 8 days of development (fig. 3). Two days after hatching most animals develop two red chromatophores anterior to the central fluorescent chromatophore and two which are positioned directly on either side along the boundary of the 2^nd^ and 3^rd^ tergites (fig. 3A-B). Overlapping red and blue pigmentation also develops on the D3 chromatophore starting at around 2 days post hatch (fig. 3B-B”) and persists throughout larval development.

**Fig. 3.**
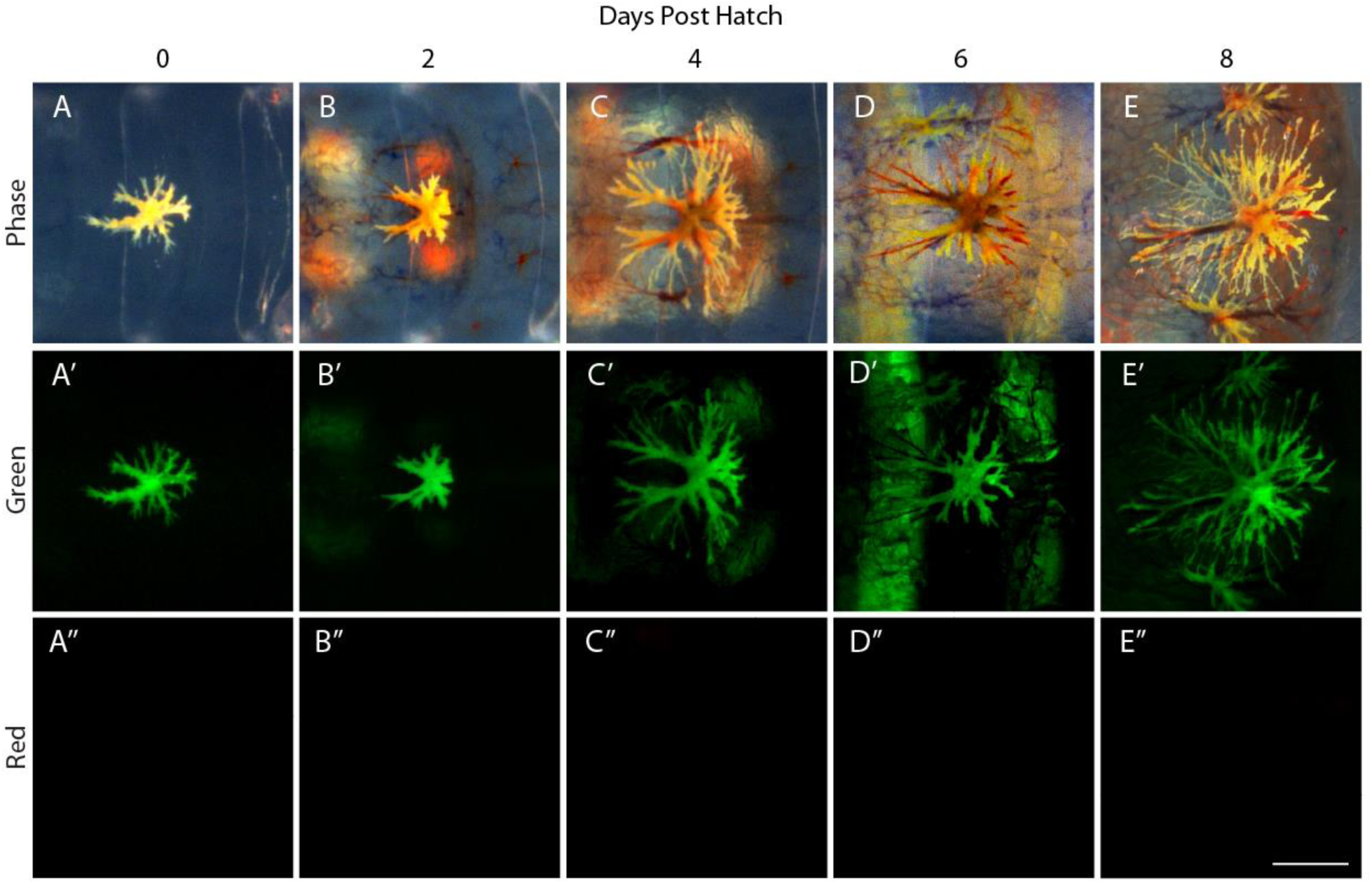
Time series images of an individual dorsal D3 chromatophore from hatch until 8 days post hatch. No red chromatophores were present at hatch (A-A’’), however these cells rapidly surround the D3 chromatophore by 2 days post hatch (B-B’’). Fluorescent pigment accumulates in or around the flanking red chromatophores by day 4 post hatch (C-C’’). This fluorescence gradually increases through day 6 (D-D’’) and day 8 (E-E’’) post hatch. The V2 chromatophores can be observed through the transparent larva as background yellow/red pigmentation and green fluorescence. Phase contrast (Phase), green (’) and red (“) fluorescent filters are identified. Scale bar represents 100μm.

Four days after hatching the lateral red chromatophores develop green fluorescent, yellow/white pigmentation (fig. 3B-B’’). The fluorescent pigmentation associated with these cells gradually increases through day 6 (fig. 3C-C”) and 8 (fig. 3D-D”) post hatch but does not appear to reach the extent of the original lone dorsal chromatophore (D3; fig. 2C-C’). While the red chromatophores located on the anterior side of chromatophore D3 are present throughout larval development they do not develop fluorescent pigmentation (fig. 3).

### Post larval fluorescent chromatophores

Green fluorescent chromatophores persist after post larval metamorphosis but there is a major restructuring of the pigmentation pattern. Instead of localized clusters and overlapping chromatophores as is found in larval organisms, the fluorescent chromatophores of the post larvae are systematically dispersed throughout the organism (fig. 4A-A’’). The fluorescent chromatophores of the head form a distinct line between the eyes which extends the length of the eye stalks (fig. 4B-C). Fluorescent pigment cells also reside on the base of the antennule flagella and the distal end of the antennule peduncle (fig. 4D-D’). While ventral fluorescent chromatophores are present in post larvae, the chromatophores do not overlap surrounding cells (fig. 4 E-E’) as is seen during larval development. A distinctive feature of post larval pigmentation that is not found prior to metamorphosis is the high concentration of fluorescent chromatophores which resides within the animal. These chromatophores appear to uniformly cover the hepatopancreas and parts of the stomach and are found in all post larval animals (fig. 4 f-f’).

**Fig. 4.**
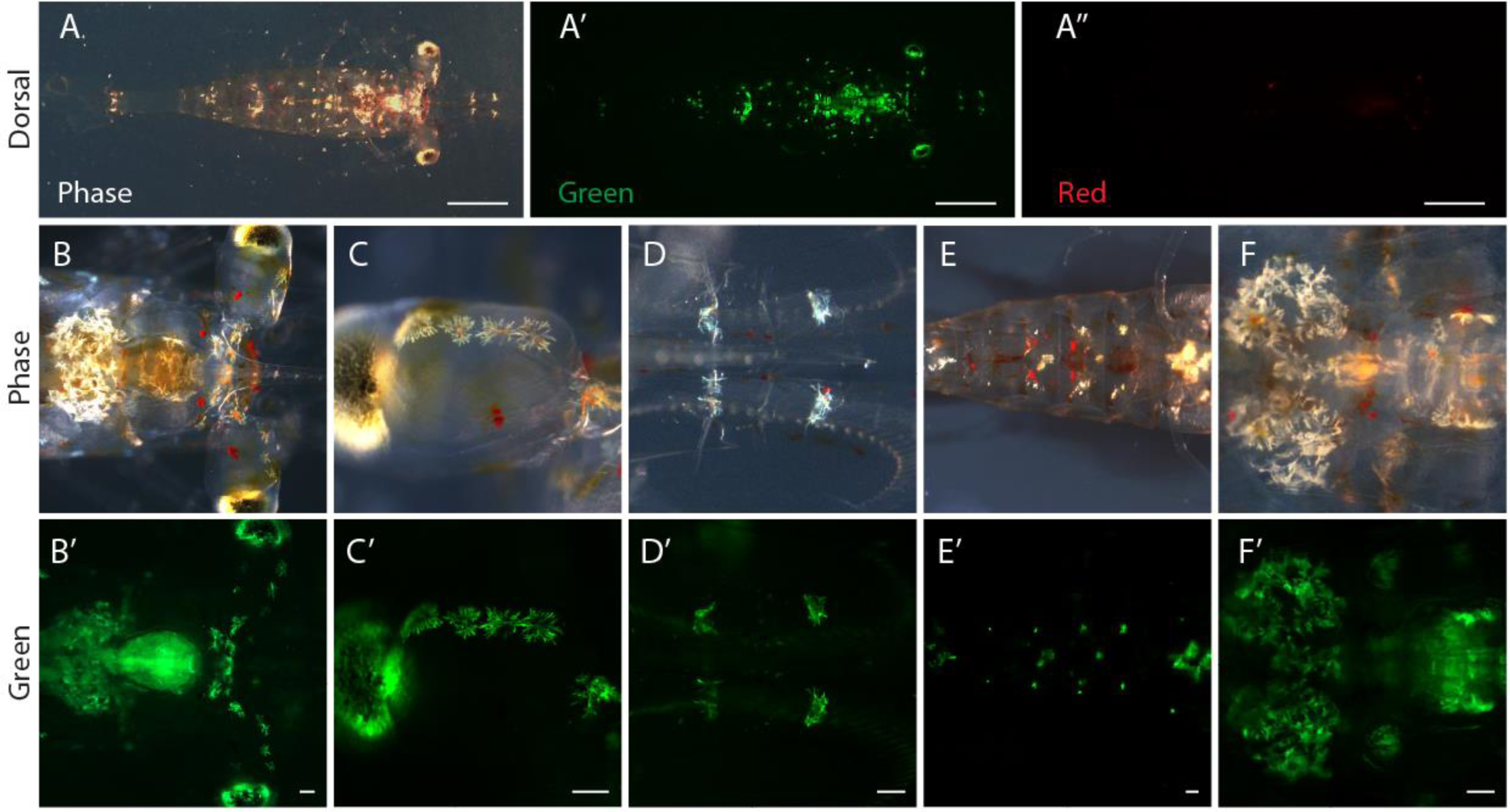
Redistribution of chromatophores in post-larvae. Green fluorescent chromatophores are more defined and broadly distributed throughout the post larvae (A-A”) and are primarily white in coloration. The highest concentration of these chromatophores appear to cover the hepatopancreas and parts of the stomach (B-B’, F-F’). There is also a prominent row of fluorescent chromatophores between the eyes (B-B’, C-C’). Pairs of individual chromatophores are present on the antennules (D-D’) and on the ventral sternites (E-E’). Image taken with phase contrast (Phase), green (’) and red (“) fluorescent filters are identified. Scale bars for whole body images (A-A”) and regional chromatophores are 1mm and 100μm, respectively.

The fluorescent chromatophores of post larvae do not resemble those of larval organisms but are more uniform in shape and have lost most of the yellow pigmentation found in larval chromatophores (fig. 4). The associated red and blue pigments that commonly overlap with larval fluorescent chromatophores are also limited in post larvae with most cells only exhibiting a small centralized patch of red pigmentation.

### Fluorescent chromatophores in adult *P. pugio*

The fluorescent pigmentation pattern of adult *P. pugio* varies significantly between individuals, to a larger degree than in larval and post-larval organisms. Individual shrimp cultured under identical conditions showed both the presence or near absence of fluorescent chromatophores (fig. 5 a-d). When green fluorescent chromatophores were present they were most commonly located on the antennules (fig. 5a-a’), distributed between the eyes (fig. 5b-b’), and on small centralized patches on the posterior margin of tergites (fig. 5c-c’). While this pigmentation pattern has similarities to that observed in post larvae, overall chromatophore abundance was markedly reduced in adults.

**Fig. 5.**
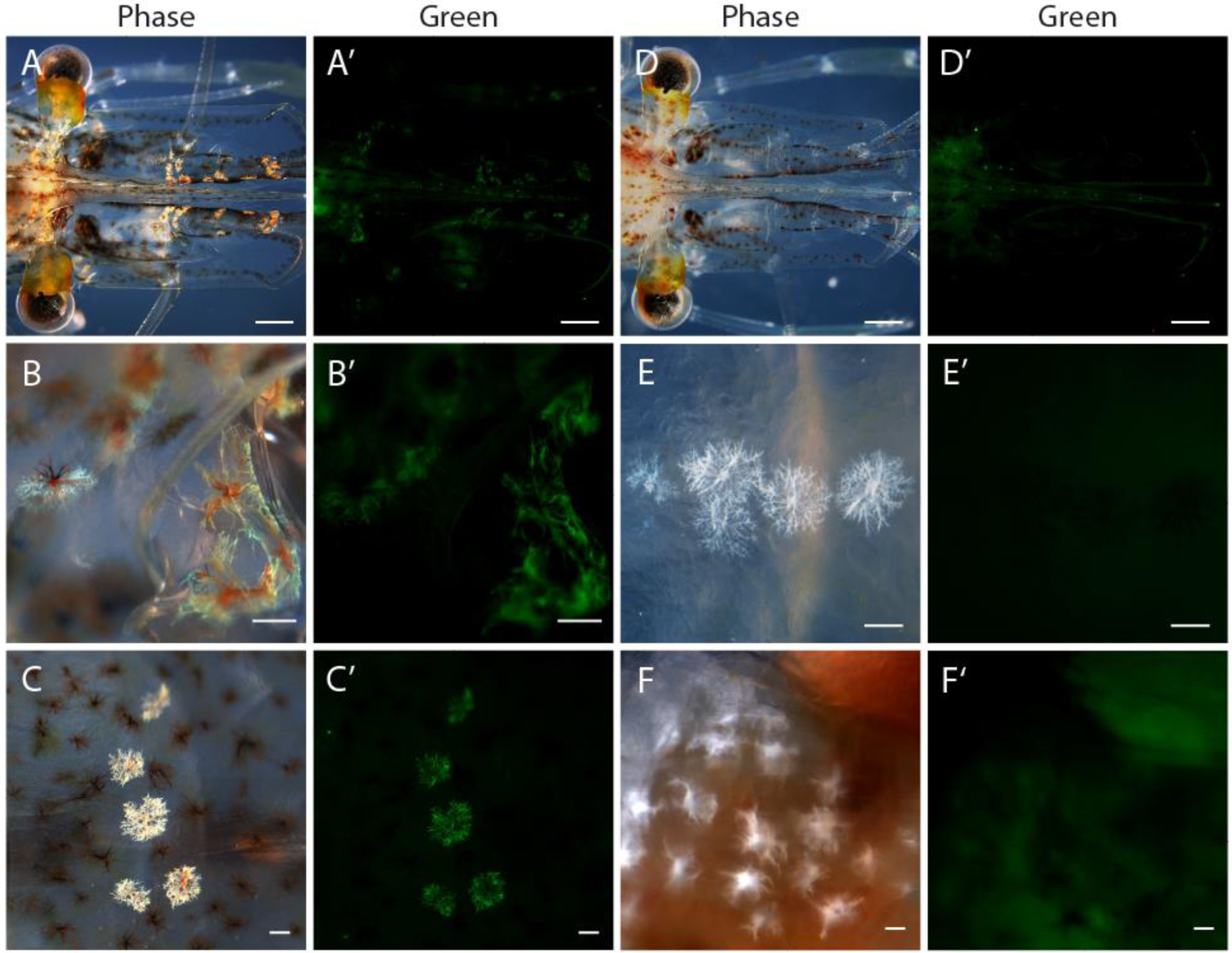
Variation in the presence of fluorescent chromatophores in adult female *P. pugio.* Some individuals retain green fluorescent chromatophores on the antennules (A-A’), between the eyes (B-B’) and patches on the dorsal tergites (C-C’). Not all individuals however exhibit these fluorescent cells (D-D’), and not all white pigmented cells are fluorescent (E-E’, F-F’). This lack of fluorescence in white pigmented cells can be found on some dorsal chromatophores (E-E’) but more prominently is a characteristic of white chromatophores found on the hepatopancreas and stomach (F-F’). Low level background autofluorescence of the internal organs and seta of adult organisms is common which can be detected in all filter channels (A-A’, D-D’, and F-F’). Images under phase contrast (Phase) and fluorescent green filter (’) are shown. Scale bars for images of the adult head (A-A”, D-D’) and regional chromatophores are 1mm and 100μm, respectively.

Adult shrimp also possessed white chromatophores that did not exhibit green fluorescence (fig. 5e-e’). This was particularly evident with chromatophores of the hepatopancreas, which were readily distinguishable in adults, yet lacked detectable green fluorescence (fig. 5f-f’).

## DISCUSSION

This study tracks the development and distribution of green fluorescent chromatophores in the grass shrimp *P. pugio*. While fluorescent chromatophores were present in all life stages, including during embryonic development, they were most prevalent in larval and juvenile shrimp. In fact, fluorescent chromatophores were the only pigmented cells present during embryonic and early larval development. The chromatophores occupied several large patches throughout the body, concealing surrounding transparent tissues. The pattern of chromatophore distribution was well conserved between individuals during larval and juvenile life stages. This was not the case for adults, which showed significant variation in the presence and distribution of fluorescent chromatophores between individuals.

The prevalence and distribution pattern of green fluorescent chromatophores in *P. pugio* raises intriguing questions about the adaptation of pigmentation in these transparent shrimps. This is especially important given the fact that the presence of pigmentation directly impedes efforts to maintain a transparent camouflage. Clues to the potential benefits of developing fluorescent chromatophores may be gained from analyzing the pattern of chromatophore distribution throughout development. It is possible that since these cells arise early in development and are prominent throughout the planktonic life stages that they may have a role in attracting prey, which could out weight any negative fitness costs associated with reducing the organism’s transparency.

Pigment cells can also have a significant role in protecting organisms from the harmful effects of UV radiation. In fact, accumulation of protective pigments such, as carotenoids or melanin and behavioral avoidance are the two primary mechanisms crustaceans employ to minimize exposure to UV radiation (Johnsen, 2001; Gouveia et al., 2004; Hansson, 2004; Dahms and Lee, 2010). In transparent species however, increasing pigmentation for UV protective purposes reduces the effectiveness of the transparent camouflage enhancing the risk for predation (Johnsen and Widder, 2001; Hansson, 2004). Previous research has shown that the chromatophores of *Palaemonetes spp.* exhibit a pigment dispersion response to UV radiation suggesting the potential involvement of these cells in UV protection (Gouveia et al., 2004). By adapting white chromatophores with fluorescent pigments or proteins, these post-larval organisms may enhance UV protection while maintaining relative transparency (i.e., no dark pigmentation). Further research is needed to investigate the potential adaptive advantage of these fluorescent pigment cells in the early life stages of *P. pugio*.

The yellow/white chromatophores exhibiting fluorescence in *P. pugio* are similar to vertebrate xanthophores which can exhibit green fluorescence due to pteridine production. Green fluorescent pteridine pigments have been documented in the chromatophores of several species including freshwater zebrafish (*Danio rerio,* Hamilton, 1822; Odenthal et al., 1996; Guyader and Jesuthasan, 2002), amphibians (Bagnara and Obika, 1965; Epperlein and Claviez, 1982; Epperlein and Löfberg, 1984), and reptiles (Ortiz and Williams-Ashman, 1963). The majority of fluorescent pteridine pigments have been identified from purified pigment extracts, which can have bright fluorescent properties (Bagnara and Obika, 1965). Pteridine containing xanthophores however mostly exhibit faint cellular fluorescence, and often require chemical treatment to enhance the fluorescent signal (Epperlein and Claviez, 1982; Epperlein and Löfberg, 1984; Odenthal et al., 1996; Guyader and Jesuthasan, 2002). This differs significantly from the fluorescence observed in the chromatophores of *P. pugio,* which have strong green fluorescence readily detectable, with up to 60 fold less exposure time than reported for similar imaging of xanthophores in vertebrates (Guyader and Jesuthasan, 2002).

Another potential source of green fluorescence, which cannot be ruled out at this time, is the production of a green fluorescent protein. Green fluorescent proteins are not uncommon in lower marine invertebrates but the vast majority of organisms exhibiting this trait are Cnidarians (Matz et al., 1999; Shagin et al., 2004; Chudakov et al., 2010). It is often the case that natural green fluorescent proteins are not associated with distinct chromatophores (Baumann et al., 2008; Hunt et al., 2010), however fluorescent proteins are present in the pigment cells of some corals (Schlichter, Meier, and Fricke, 1994; Salih et al., 2000). While the only Bilaterians that have been found to possess green fluorescent proteins are specific species of copepods (Shagin et al., 2004) and the lancelet *Branchiostoma floridae* Hubbs, 1922 (Baumann et al., 2008; Bomati, Manning, and Deheyn, 2009), non-fluorescent members of this protein family are common among metazoans (Shagin et al., 2004; Bomati, Manning, and Deheyn, 2009).

The discovery of fluorescent chromatophores in the grass shrimp *P. pugio* stimulates many questions about the adaptation of pigmentation in this transparent shrimp. This study provides a comprehensive analysis of the development of these unique fluorescent chromatophores tracking the expansion and movement of these cells throughout the life stages of *P. pugio*. How these fluorescent chromatophores play a role in the life history of grass shrimp and whether they are conserved in other transparent invertebrates could have important implication for our understanding of the ecology of shallow water coastal ecosystems.

## ACKNOWLEDGEMENTS

The microscopy resources used in this research were generously provided by Zipora Yablonka-Reuveni at the University of Washington. The author would also like to acknowledge Ian Jaffe and Terrence Bradley for contributions to reviewing the manuscript. All research, data interpretation and manuscript writing were completed by M.P.

